# Endometrial microbiome in mares with and without clinical endometritis

**DOI:** 10.1101/2025.03.06.640270

**Authors:** Lulu Guo, G. Reed Holyoak, Udaya DeSilva

## Abstract

Chronic endometritis (CE) has been recognized as an important disease in the clinical theriogenology practice. Studies have suggested that CE is one of the major causes of infertility in breeding mares. However, a comprehensive analysis of the CE microbiome is currently insufficient in published reports. In this paper, we compared the uterine microbiomes of mares with CE to those of healthy mares and propose that there are significant differences in the composition of the uterine microbiome between these two groups of mares. This study suggests that changes in the uterine microbiome may play a vital role in the development and progression of CE in mares, adds to the understanding of the role of the uterine microbiome in CE and in developing targeted treatment strategies.

## Introduction

There is strong evidence that chronic endometritis (CE) is an important factor causing female reproductive failure. CE destroys the commensal coexistence between microorganisms and the host immune system in the uterus (Kim et al. 2016). Chronic inflammation creates a hostile environment that may either be embryotoxic, and/or hinders normal placentation of an embryo and its subsequent development. It occurs in various animal species, including humans and mammals such as cows, mares, and bitches. Chronic endometritis refers to an extended duration of inflammation of the endometrium. It can lead to reproductive issues and infertility in affected humans and animals. Diagnosed CE was detected in 30% of infertile women (Cicinelli et al. 2005), 18.1 % of infertile cows (Shabunin et al. 2020), 15% in bitches over 8 years of age (Asa et al. 2014) and from 25 to 60% in infertile mares (Hurtgen 2006; Riddle et al. 2007; LeBlanc & Causey 2009). One study of Thoroughbred mares in the United Kingdom reported that 34% of mares diagnosed with infertility had CE (Pycock & Newcombe 1996). Likewise of Warmblood mares in Germany 32% of infertile mares had CE, and that the severity of the disease correlated with decreased pregnancy rates (Kohne et al. 2020). The American Association of Equine Practitioners (AAEP) lists CE as one of the most common causes of infertility in mares in North America.

Therefore, CE is a leading factor of infertility or reduced fertility in breeding mares and can have a marked economic impact on the equine industry. It can lead to reduced conception rates and an increased number of cycles and treatments required to achieve pregnancy, which result in additional expenses associated with repeated breeding attempts, including veterinary fees, treatment costs, semen collection and processing costs, and transportation expenses. It also results in lower foal production rates, which can directly impact the profitability of breeding operations. Therefore prevention, early diagnosis and appropriate treatment are very important.

The diagnosis of CE is hampered due to there being no pathognomonic clinical signs or ultrasound findings. In women, it is based on the identification of endometrial stroma’s plasma cells, classic CE diagnostic methods depend on histology (Greenwood & Moran 1981), but this method is non-specific and dependent on the menstrual cycle’s date when sampling proceeds. In the mare, while endometrial cytology and microbial culture are most commonly used for CE diagnosis (Cicinelli *et al*. 2005; Cicinelli et al. 2009; Katila 2016; Nocera et al. 2021), the most common clinical sign is the presence of luminal fluid within the uterus (LeBlanc 2010). But that is subjective because the normal equine estrous cycle induces endometrial edema and mild luminal fluid accumulation. Microbial culture identification of endometrial pathogens is the only way to provide objective information for targeted therapy. However, endometrial bacterial culture is not always useful since it often has a relatively long turnaround time, and likely not all microorganisms leading to CE are culturable. The uterine microbiome refers to the diverse community of microbiota that inhabit the uterus. Therefore, we looked at the microbiome of a group of clinically healthy mares and those with diagnosed CE. Here we provide some preliminary findings of the uterine microbiomes between the healthy mare and symptomatic mare by applying high throughput sequencing technologies to add insight into diagnosing and possible treatments for CE.

To our knowledge to-date, the comprehensive microbiome analysis in CE horses is not well established. These preliminary findings begin the development of an essential data bank.

## Methods

### Experimental design and sample collection

A total of 26 mares were selected for this study. Mares ranged from 4 to 18⍰years of age and did not have recent antibiotic exposure. All the enrolled mares are from Oklahoma, but different ranches. Five of CE and five healthy mares were from a facility in South-central OK. Four samples in each group were from a facility in central OK, and the rest of the samples were from mares from the Oklahoma State University Veterinary Medicine Ranch. These mares were long-term occupants at their respective resident facilities with similar diets. In total, 13 healthy mares with normal reproductive histories and no clinical signs were recruited as the healthy group were matched by location and diet to 13 mares previously diagnosed with CE and assigned to CE group in this study.

Endometrial microbial lavage samples were collected with a sterile triple-guarded system to minimize the risk of vaginal microbial contamination as described in equine microbiome studies (Holyoak et al. 2022). The perineum and external genitalia were cleaned 3 times with an iodophor scrub and alcohol to minimize external contamination. Sterile obstetrical lubricant was used on a sterile disposable speculum before entering the reproductive tract. A sterile obstetrical sleeve was applied to protect and guide the speculum and catheter through the cervix. 150ml of saline was infused, agitated within the uterine lumen, and after 30s of contact with the endometrium, 45ml of lavage was decanted and collected midstream. All samples were stored on ice temporarily and transported to the laboratory in no more than 2h. The samples were processed in the laboratory immediately within 2h after sampling.

### Negative Control

Negative controls were collected by passing through the same system used in regular sample collection but without putting into the reproductive tract in all barns when collecting mare samples. DNA isolation and 16s rRNA amplification were processed with the two sample groups together to avoid potential unnecessary impact on external conditions caused by human factors. All the Laboratory reagents were the same. Sample treatment and subsequent sequence analysis were performed by a single investigator. Totally, three negative control samples were collected and used in this study. All negative control samples failed in amplification and sequencing steps.

### DNA extraction

45 mL of lavage for each sample was spun down at 7000 rpm for 15 min at 4 °C to settle down the suspending bacteria. DNA of all CE, healthy as well as negative control samples were extracted in a controlled environment by using a QiAamp DNA MINI KIT (Qiagen, Germantown, MD) depending on manufacturer’s instructions. Quality and quantity of DNA products were measured with spectrophotometer (NanoDrop 1000, Thermo Scientific, Wilmington, United States).

### 16S rRNA gene amplicon sequencing

The PCR amplification of extracted DNA and 16S amplicon sequencing was conducted by universal primer set of 515FF (5′-GTGCCAGCMGCCGCGGTAA-3′) and 806R (5′-GGACTACHVGGGTWTCTAAT-3′), which targets the bacterial V4 hypervariable region (∼250 bp) by using Phusion^®^ High-Fidelity PCR Master Mix (New England Biolabs). Barcode sequences were carried in forward primer.

The PCR products were verified by 1.5% agarose gel electrophoresis and extracted using a Qiagen^®^ Gel Extraction Kit (Qiagen, Germantown, MD). NEBNext^®^ UltraTM^®^ DNA Library Prep Kit (New England Biolabs, Ipswich, MA) were used in library preparation for Illumina. Qubit 3.0 fluorometer and Q-PCR were used for qualification and quantitation, respectively. Subsequently, an Illumina HiSeq 2500 platform was utilized for the sequencing of the library. Purified amplicons were pooled and generated into paired-end sequence reads on an Illumina HiSeq 2500 platform (Illumina, San Diego, United States).

### Bioinformatic and Statistical analysis

The sequences were mainly analyzed using the latest QIIME2 (Hall & Beiko 2018). Data preprocessing is based on the overlap between reads. The paired-end sequence data obtained by HiSeq sequencing were merged to sequence tags. Then quality control filtering was performed on detecting the reads quality and the merge effect. There are mainly three steps. First, the reads of each sample were merged by overlapping, and the merged sequence obtained was the original tags data (Raw Tags). Secondly, filtered the merged raw tags to obtain high-quality tags data. Lastly remove chimeras, by identifying and removing chimera sequences to obtain final valid data.

UCLUST in the QIIME2 was used to cluster the tags at a similarity level of 97% to obtain OTUs, and the OTUs were taxonomically annotated based on the Silva (bacteria) and UNITE (fungus) taxonomic databases. In this step of QIIME2, there are mainly two methods, DADA2 (Babtiwale et al. 2021) and Deblur (Amir et al. 2017). According to (Callahan et al. 2016) in Nature Method, when comparing similar methods DADA2 is better than other OTU clustering results so we chose DADA2. Compared with QIIME’s UPARSE clustering method, the current DADA2 method will only remove noise and chimera but cluster by similarity. It has more accurate analytic results than the previous generation. The main function of DADA2 is to remove low-quality sequences and chimeras and regenerate the OTU table, which is now called the feature table. Because the clustering method is no longer used, the feature table is equivalent to the OTU table with 100% similarity in the QIIME era.

All statistical parts were completed in R language. Alpha diversity was analyzed using Shannon indexes, and student’s t-test was performed to illustrate the differences of the indexes between the two groups. Differences in community composition were demonstrated by Wilcoxon Test. The beta diversity was analyzed by non-metric multi-dimensional scaling (NMDS) using the ape software package in R language, and the differences between groups and within groups were evaluated by the analysis of similarity (ANOSIM) statistical test. Linear discriminant analysis (LDA) effect size was applied to identify the differentiated bacteria between 2 groups. Tax4Fun software was utilized to infer the functional gene composition in the sample by comparing the species composition information obtained from the 16S sequencing data, so as to analyze the functional differences between different groups.

## Results

Negative control samples for both CE and healthy samples failed in two-step PCR amplification protocol for 16s rRNA V4 region, which indicates that bacteria in the negative control samples were negligible (Siddiqui et al. 2011). Rarefaction curves performed on CE and healthy groups indicated that the sequencing depth were sufficient to cover the overall bacterial diversity in all samples. A total of 1,913,005 sequence reads were detected from the 13 healthy mare samples, and 5,868,329 sequence reads were retained after removing the host and low-quality reads. More than 90% of reads passed denoise and chimera removal. Totally, 11182 OTUs were obtained. 33621 OTUs in CE mare group and 22439 OTUs in healthy mare group, were yielded to downstream analysis.

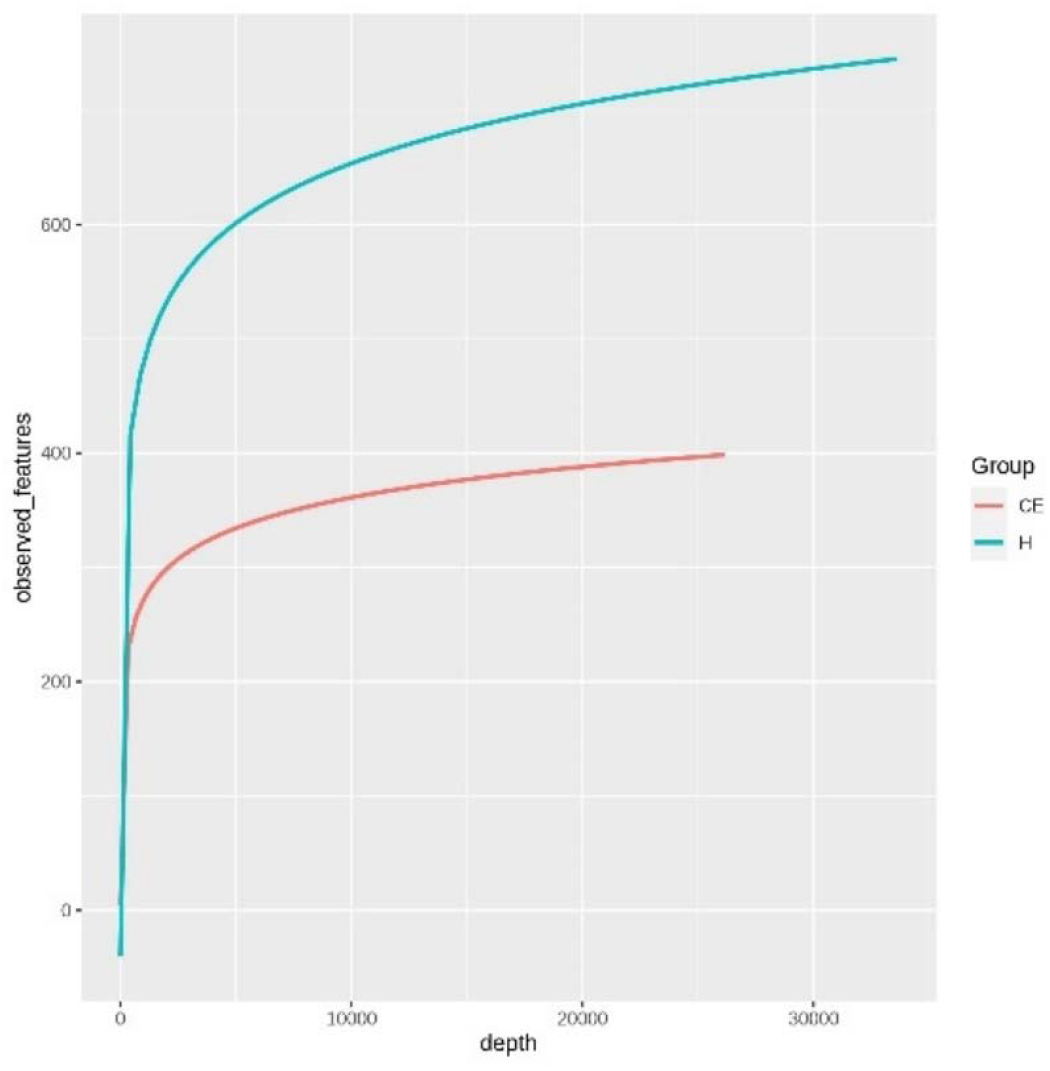

Among the microbiome differences between the heathy group and the CE group, *Burkholderia* and *Chlamydia* were the most abundant genus in the endometrial microbiomes of both groups. However, Wilcoxon Test showed Burkholderia (p-value = 0.024665⍰<⍰0.05) and Chlamydia (p-value = 0.05 ⍰≤ 0.05) in the CE group had significantly higher relative abundance than the healthy group. The rest of the genera with lower relative abundance were *Pseudomonas, Enterobacteriaceae* and *Buchnera*. There was a significant difference (p-value = 0.00262⍰<⍰0.05) between the two groups based on the top 50 genera by Wilcoxon Test. Venn diagram demonstrated that 608 genera were shared by the two groups. 292 and 211 exclusive genera were identified in the healthy group and CE group, respectively. The heatmap of the top 50 genera showed the CE group had decreased clustering compared to the healthy group.

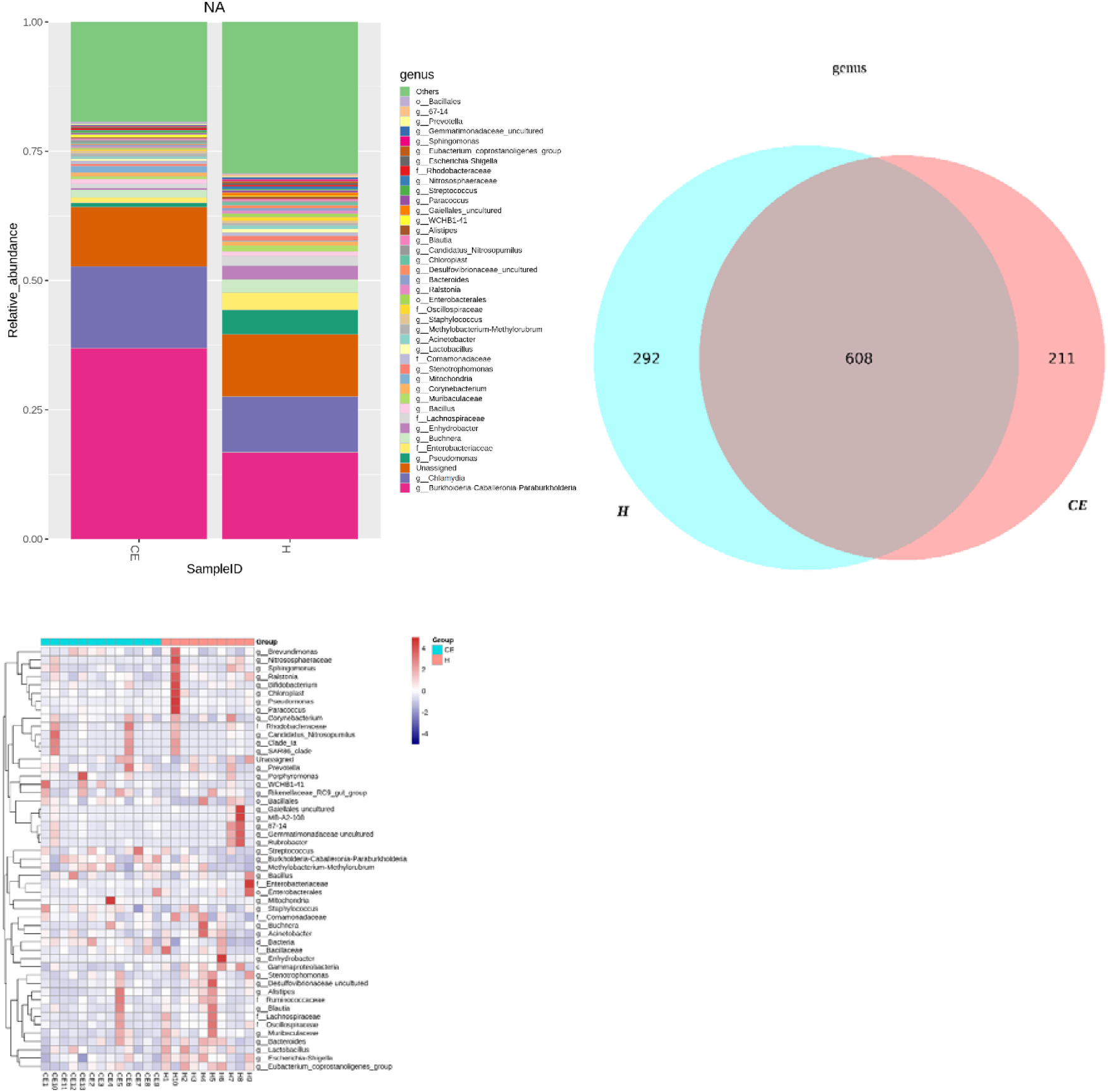

The alpha diversity in the Shannon index was used to calculate the richness and evenness in the microbiome and indicated the significant differences between 2 groups via t-test (p-value = 0.049 <⍰0.05). The healthy group was statistically higher in richness and evenness compared to the CE group.

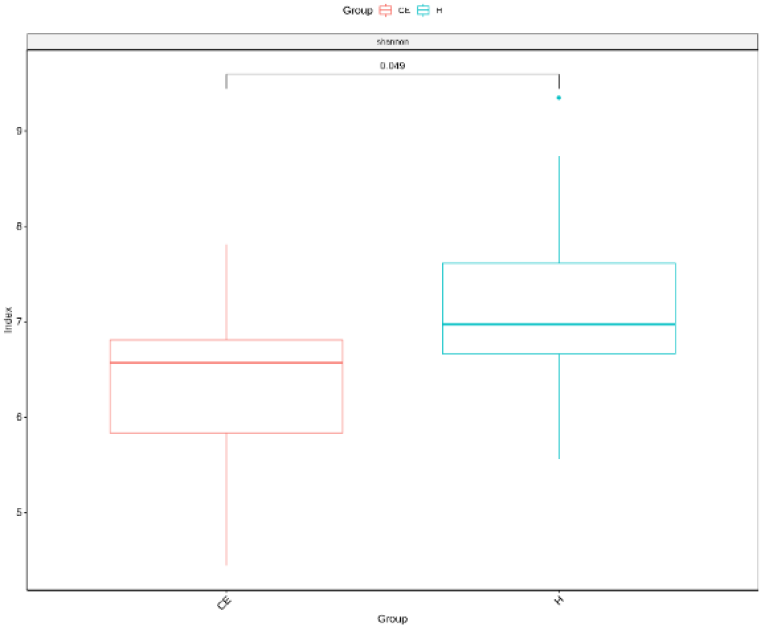

Unlike the alpha diversity analysis, beta diversity is used to reflect the similarity or dissimilarity of the microbial communities between groups. Principal coordinate analysis (PCoA) with Bray-Curtis dissimilarity showed two distinct clusters about community composition.

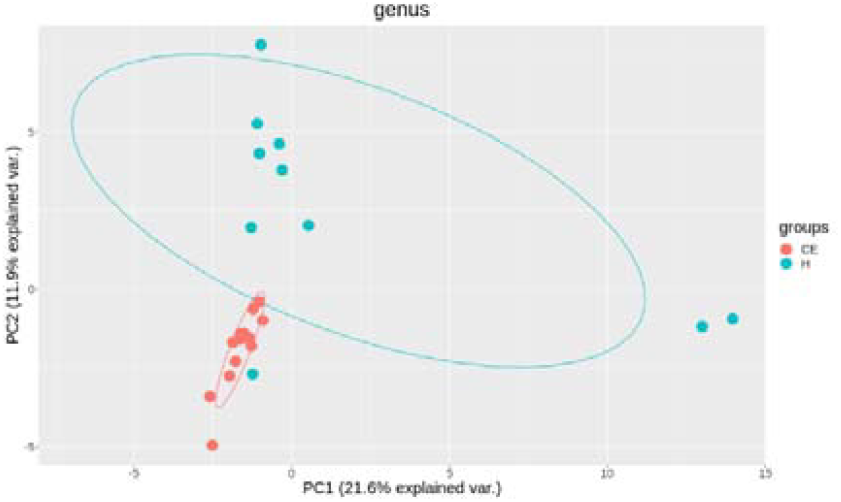

A petal diagram was drawn to analyze the common and unique information of microbial diversity between the groups. The central circle of the petal diagram represents the number of phyla shared by all samples, and the numbers on the petals represent the number of phyla unique to the to the group of samples. The unique phyla found only in the CE group are Halobacterota, uncultured delta proteobacterium Sva0485 and WPS-2. WPS-2 is a phylum that is evolutionarily near to Chloroflexi,

Armatimonadetes and Dormibacteria. The unique phyla in the healthy group were Acetothermia, Caldatribacteriota, Deferribacterota, Margulisbacteria and Fibrobacterota.

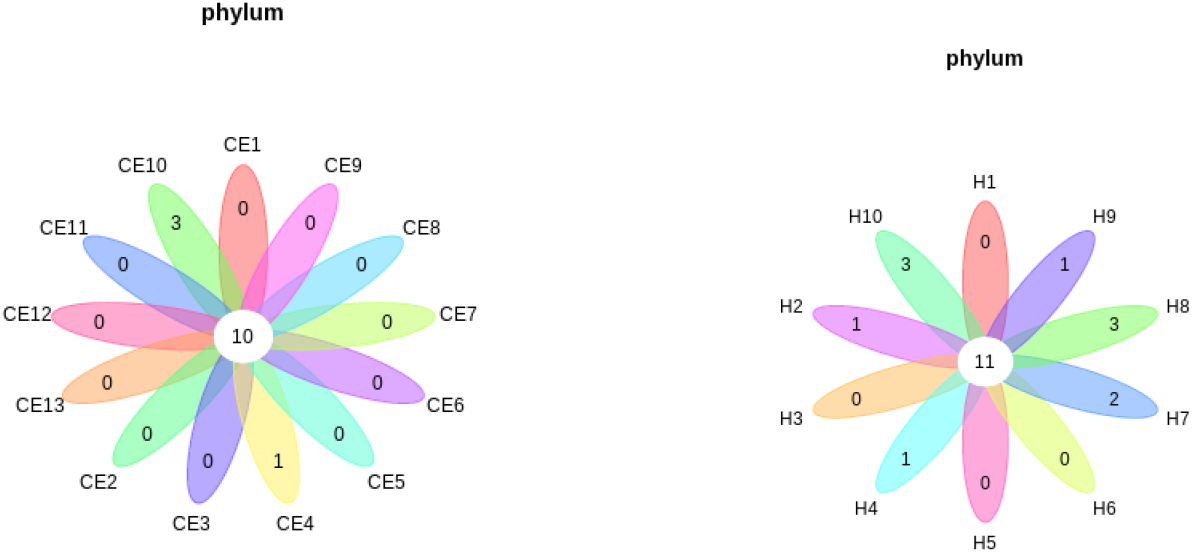

To filter species with significant differences between groups Biomarker, LEfSe with a linear discriminant analysis score⍰>⍰2 was applied to confirm the differentially abundant taxa. Burkholderia (genus), Burkholderiales (order), Hyphomicrobium (genus), and Erwiniaceae (family) had significant association with CE mares with higher LDA scores.

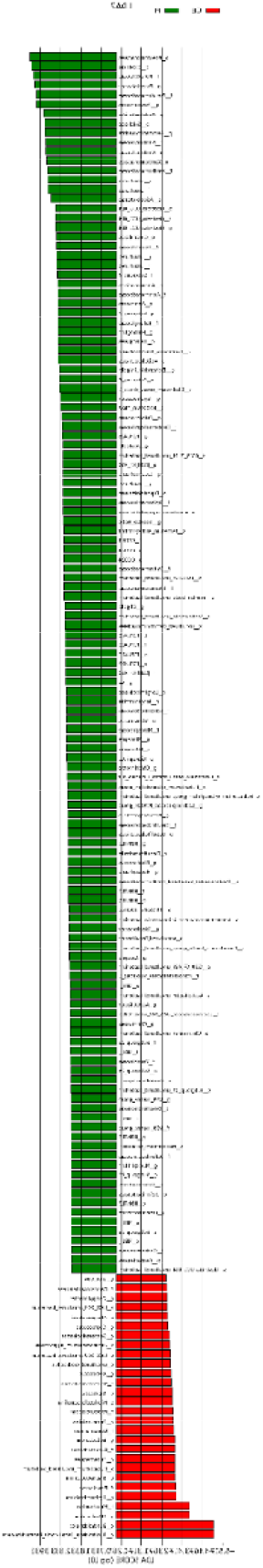

To study the species with significant differences between groups, the Metastats method was used to perform hypothesis testing on 7 different abundance level data between groups to obtain the p-value, and to screen the species with significant differences according to the p-value species. Boxplot of the abundance distribution of different species between groups indicated Erwiniaceae at the family level, Anoxybacillus at the genus level and 2 other uncultured species were significantly higher in the CE group than the healthy group. Clostridium species and Roseburia genus were richer in the healthy mares.

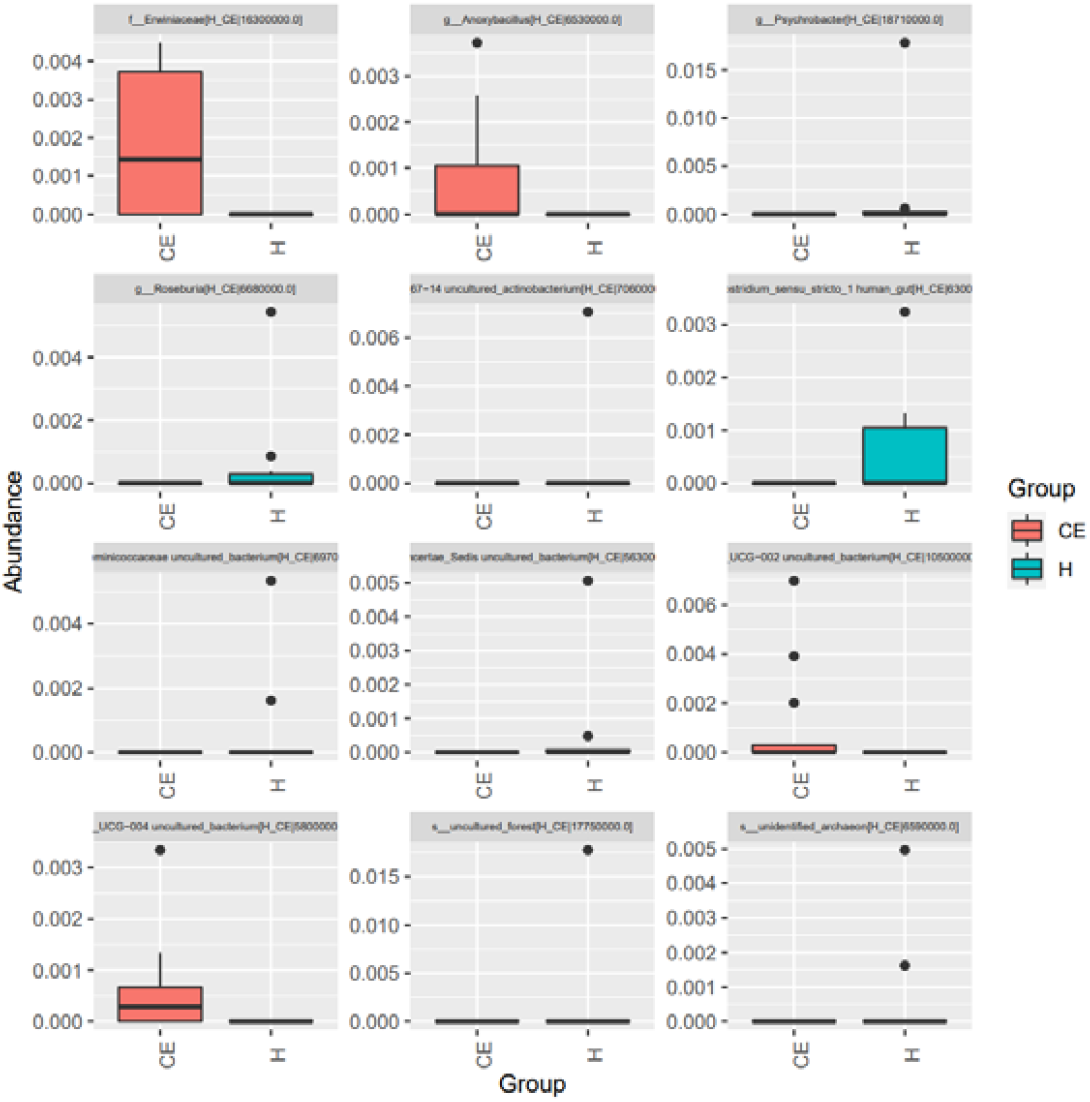

The functional profile of the endometrial microbiome was compared between the CE and healthy groups by KEGG pathway analyses. Welch’s t-test with 95% confidence intervals were used through STAMP. Microbiota in the CE group showed significantly less richness than the healthy group in pathways of Brite Hierarchies factors. In addition, metabolism was significantly more abundant in the CE group compared to the healthy group.

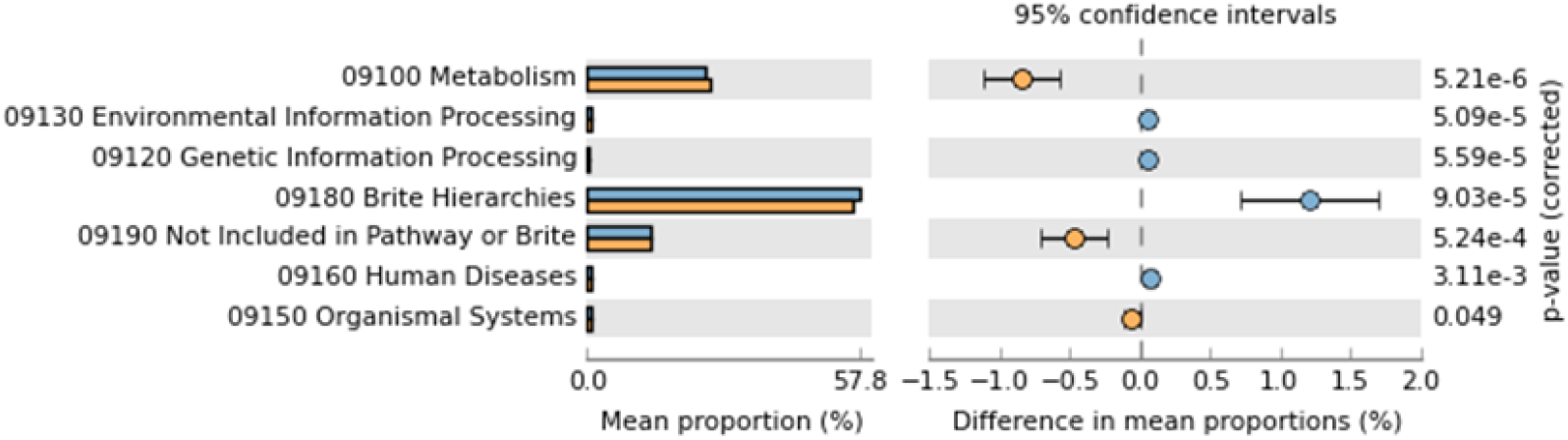

## Discussion

There is growing interest in the role of the mare uterine microbiome in the development and progression of chronic endometritis. Research in this area is needed to lead to improved diagnosis and treatment strategies for CE in mares. This study has investigated the differences in the uterine microbiome between relatively small groups of mares with CE and clinically healthy mares in order to better understand the potential microbial interactions and causes of the disease.

Although the most abundant microbial compositions of the two groups were similar, the ratio of bacteria were different. The composition in the healthy group was obviously richer than the CE group, which indicates that the diversity of the microbiome helps maintain local health. In healthy mares, the uterine microbiome is typically high in diversity, which helps to maintain a healthy environment in the uterus. The imbalance of certain bacteria may consume local resources and otherwise suppress resident microbes, thereby decreasing the whole diversity of the microbiomes and inducing dysbiosis. These observations suggest that dysbiosis is the pathogenesis of CE.

Burkholderia-caballeronia-paraburkholderia was the most abundant genus in the CE group and Burkholderia species can include pathogenic strains. However, it is important to note that Burkholderia, Caballeronia, and Paraburkholderia are three genera of bacteria within the Burkholderiaceae family. These genera are closely related and share similarities in their genetic and phenotypic characteristics. They should be further reclassified based on genomic and phylogenetic analysis to better reflect their evolutionary relationships. Since there is insufficient data that could be used in the uterine region, most of the sequences are unable to be classified as accurate genera. We expect this study to contribute to reclassifying these bacteria into different genera reflecting advances in molecular techniques and in better understanding their evolutionary relationships. Reclassification will help in delineating the broadly diverse Burkholderiaceae family and providing a more accurate taxonomy for these bacteria.

Chlamydia is a type of bacteria that can cause infections in various species. Chlamydia species are known to cause reproductive tract infections in some animals, such as cats, dogs, and ruminants (Kaushic et al. 1998; Kaltenboeck et al. 2005; Holst et al. 2010), but its involvement in chronic endometritis is not well-documented. Chlamydia infections in horses can occur, but they are usually associated with respiratory or ocular diseases (Moorthy & Spradbrow 1978; Baumann et al. 2020). This study could draw attention to Chlamydia’s role in the reproductive tract either as a commensal or pathobiont.

It is interesting that Streptococcus species and Escherichia coli are often assumed to have higher levels of pathogenic bacteria in mares with CE. It is probably because routine cultural methods are used to find the targeted bacteria in CE. However, with next generation sequencing, we obtain relevant information on the many unculturable bacteria and clearer insights on the various roles in dysbiosis verses normobiosis.

Overall, these findings clearly suggest that dysbiosis, or an imbalance in the uterine microbiome, is a contributing factor in the development of CE in mares. To better understand the complex interactions between the mare uterine microbiome and the development of CE will help in developing new diagnostic and therapeutic approaches for the disease. This work made a preliminary analysis of the differences between CE mares and healthy individuals. It will enable researchers to study and characterize these bacteria more effectively, including their pathogenicity, ecological roles, and potential mechanisms in CE. Prevention of CE is critical, but how to avoid the risk factors that may lead to dysbiosis and to achieve optimal reproductive health outcomes still require extensive research. This paper summarized the relationships in microbiome between a group of CE mares and healthy mares and aimed to provide initial information on the differences.

